# Effect of Chronic Stress on Whole Blood Transcriptome: A Meta-Analysis of Publicly Available Datasets from Rodent Models

**DOI:** 10.1101/2025.05.30.657043

**Authors:** Elizabeth I. Flandreau, Duy Manh Nguyen, Megan H. Hagenauer, Minh Nguyen, Hannah Kim, Toni Duan, Anne Bader, Stanley Watson, Huda Akil

## Abstract

**Background:** Chronic stress increases risk for neuropsychiatric disorders in humans. By modeling stress-induced changes in animals, we may improve diagnosis or treatment of these disorders. Successful translation benefits from studies with sufficient statistical power and outcome measurements that can be directly compared across species. We performed a meta-analysis to examine the impact of chronic stress on the whole blood transcriptome.

**Methods:** Datasets were systematically identified in Gemma, a database of reprocessed public transcriptional profiling studies; datasets GSE68076, GSE72262, and GSE84185 met inclusion/exclusion parameters. Each study exposed eight-week old mice to chronic stress (5-10 days social defeat stress or 6-8 weeks chronic mild stress). The final sample size was *n*=92 (*n*=45 Non-Stress/*n*=47 Stress). Stress-related differential expression in each dataset was quantified using the Limma pipeline followed by empirical Bayes moderation. For the 9,219 genes represented in all three datasets, we ran a meta-analysis of Log(2) Fold Changes using a random effects model and corrected for false discovery rate (FDR). Functional patterns were assessed with fast Gene Set Enrichment Analysis. Cell type specific enrichment for each of the differentially expressed genes was further explored using a public 10x genomics scRNA-Seq dataset from mouse peripheral blood mononuclear cells.

**Results:** Findings included 23 downregulated and 16 upregulated transcripts in stress-exposed mice (FDR<0.05). Results indicated a down-regulation in gene sets related to B cells, immune response, DNA and chromatin regulation, ribosomal activity, translation, and catabolic cellular processes. Upregulated gene sets related to erythrocytes and oxygen binding.

**Conclusion:** Our results provide molecular insight into stress-related immune dysregulation and add weight to the hypothesis that environmental stress escalates cellular aging, supporting the use of blood transcriptome as a bridge between human and rodent models.

**Graphical Abstract:** 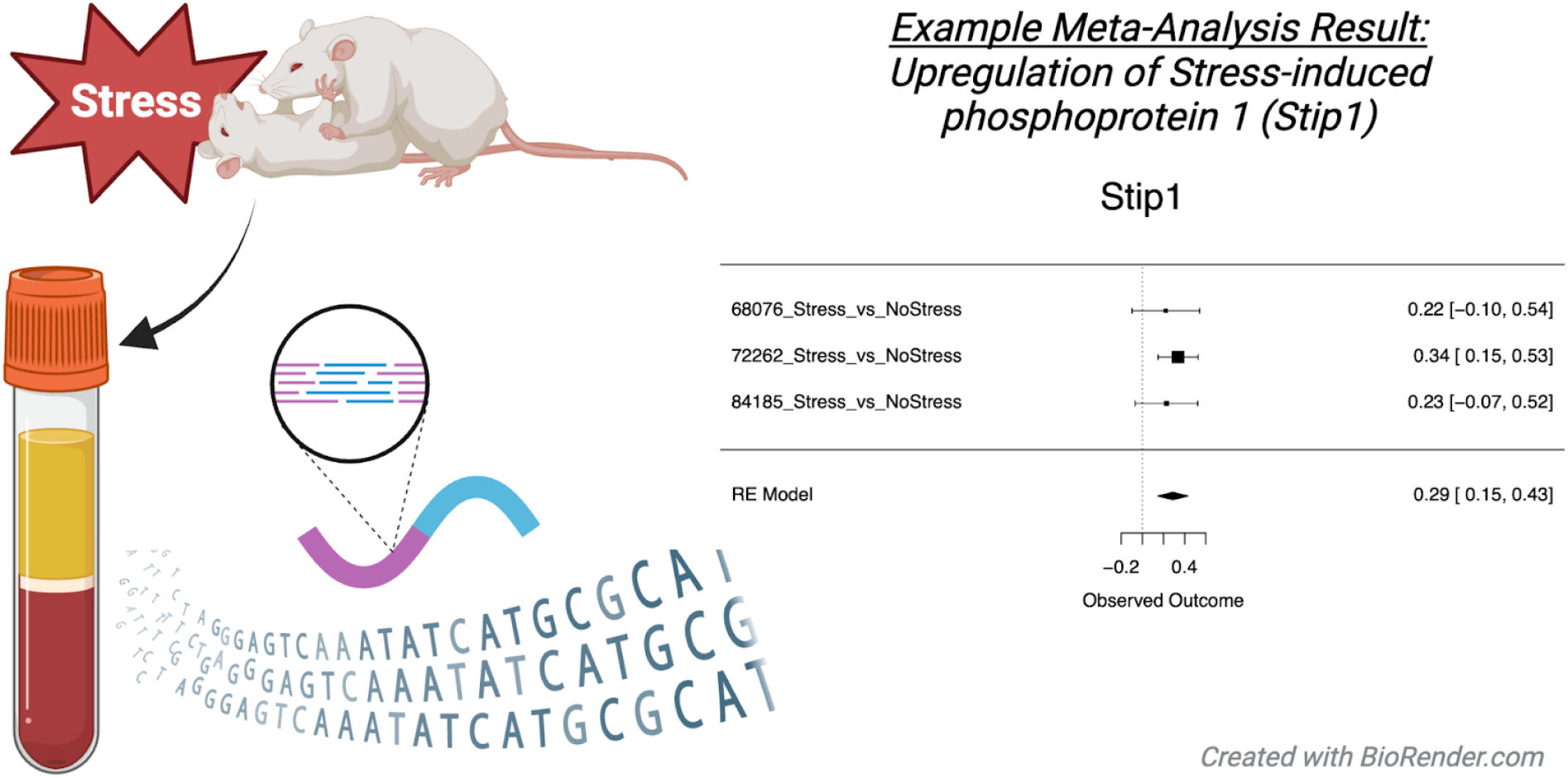

**Key points:** Successful translational research benefits from studies with outcome measurements that can be directly compared across species and sufficient statistical power. The present report is a meta-analysis of three mouse experiments examining the impact of chronic stress on the whole blood transcriptome. Our results provide insight into stress-related immune dysregulation and add weight to the hypothesis that environmental stress escalates cellular aging. These findings illustrate the utility of the blood transcriptome as a bridge between human and rodent models.

## Introduction

Exposure to environmental stress increases risk for many neuropsychiatric disorders, including major depressive disorder, generalized anxiety disorder, and post-traumatic stress disorder. Psychiatric disorders are notoriously difficult to diagnose and treat due to their myriad symptoms, heterogenous presentation, lack of objective biomarkers, and difficulty accessing the brain cells and circuits driving symptom states ^1^. These cells and circuits can be accessed in rodent models, revealing the signaling molecules and neuroanatomical pathways underlying healthy and pathological responses to stress. Yet the successful translation of these results to improvements in the clinic are few ^2,3^.

The translation of stress-related findings from animal models to humans may be improved with directly analogous measurements across species. Comparative transcriptomics has boosted translation in other biomedical fields ^4^, but brain tissue is inaccessible in human patients. Blood is accessible, however, and some brain-based transcripts can be directly measured in blood thanks to exosomes ^5^ and fluid exchange between brain and peripheral circulation ^6^. Indeed, the peripheral blood transcriptome has proven relevant to depression symptoms, providing insight into postmortem brain data ^7,8^. This suggests that the blood transcriptome in animal models of stress could provide translatable insight into psychiatric disorders.

Successful translation may also be achieved with improvements in the generalizability and reliability of results from animal models. Given the impossibility of modeling heterogeneous disorders diagnosed by patient interview, strategies to induce and evaluate “depression-” and “anxiety-” like symptoms in animals are diverse and only roughly map onto human psychiatric symptoms ^9^. Methods of stress-exposure in animal models vary greatly in duration (acute vs. chronic), modality (physical vs. psychological), and associated behavioral analysis ^10^.

The present report improves generalizability by using a meta-analysis to identify converging effects on the blood transcriptome produced by two well-established models of chronic stress: chronic social defeat stress (SDS ^11^) and chronic mild stress (CMS ^12,13^). In SDS, the experimental rodent is defeated by a larger, aggressive territorial male following 5-15 minutes of direct contact, and then spends the day experiencing additional indirect intimidation. This process is repeated for 4-10 days with different aggressors. In CMS, subjects are exposed to physical stressors (*e*.*g*., wet cage, cage tilt, altered light/dark cycle) and social stressors (overcrowding, isolation), presented in an unpredictable order over 6-8 weeks. SDS and CMS are considered valid models of stress-sensitive psychiatric disorders, increasing anxiety- and depression-like behavior in many subjects ^10^. Examining the converging effects produced by these two models may provide greater insight than either model alone.

Moreover, meta-analyses can improve power to detect reliable, true effects despite small sample sizes and methodological variability in individual datasets^14,15^S. To address the replication crisis, human ‘omics’ research is shifting from small studies in homogeneous populations toward inter-institute collaborative consortia with large, diverse datasets and consistent methodologies ^16,17^. Animal research, however, typically still relies on individual laboratory research with small sample sizes. Low statistical power ^18^ and lurking “hidden variables” ^19^ mean that false positive results may be the rule rather than the exception.

The present study aims to improve the translation of stress-related findings from animal models by performing a meta-analysis whole-blood transcriptome from mice exposed to chronic stress as a model of stress-sensitive psychiatric disorders. Combining multiple smaller studies improves statistical power as well as sample diversity in terms of sex, strain, and stress modality, increasing the probability of detecting true stress-induced differences in gene expression.

## Methods

All analyses were conducted in R Studio (R v.4.2.1, R Studio v.2023.06.0, code release: ^20,21^). Analysis followed a standardized protocol (^*22*^, *publication:* ^*23*^, *not pre-registered)*.

### Dataset Selection

Relevant datasets from rats or mice were identified in *Gemma*, a comprehensive, curated database containing nearly 20,000 reprocessed transcriptional profiling datasets ^24,25^ (function: *search_datasets()*, package: *gemma*.*R* v.1.3.2). Keywords were prespecified **(Figure 1)**. Dataset metadata and associated publication(s) were then evaluated using predefined inclusion criteria: (1) gene expression data is publicly accessible, (2) included chronic stress exposure in adult rats (*Rattus norvegicus*) or mice (*Mus musculus*), (3) transcriptome measured from whole blood using (4) RNA-seq or microarray. Prespecified exclusion criteria included: (1) flagged “problematic” by Gemma, (2) measurements reflect single cell types or cell culture or (3) a restricted subset of the transcriptome (*e*.*g*., Chip-Seq, miRNA, or TRAP-Seq). Dataset search and filtering was conducted by the first author; final inclusion decisions were approved by the full team.

**Figure 1.**
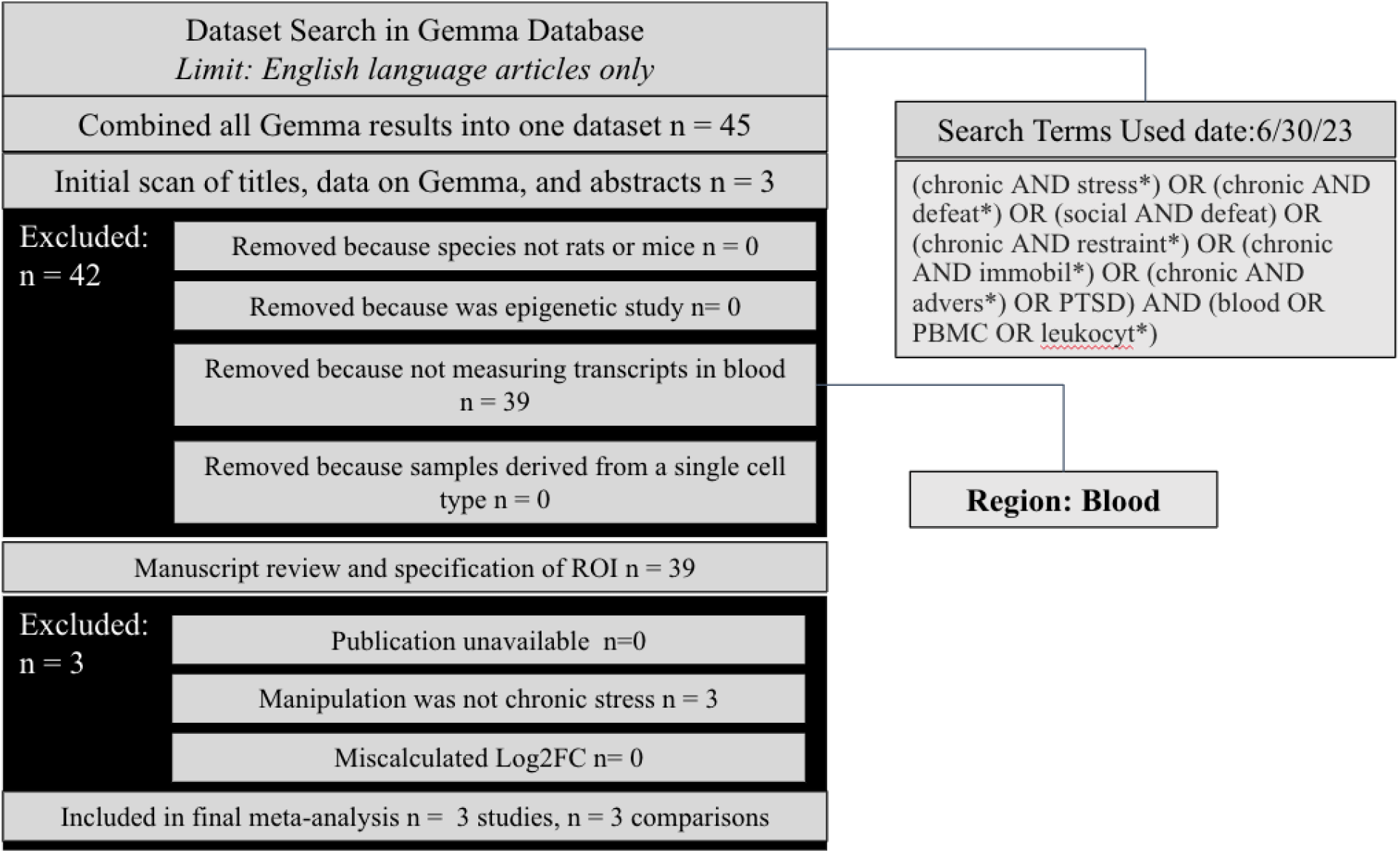
PRISMA Diagram Overviewing Dataset Search and Selection. We identified transcriptional profiling datasets examining chronic stress models in laboratory rodents within the Gemma database using pre-specified keywords. The titles, abstracts, and metadata for the datasets were initially scanned and filtered using pre-specified inclusion/exclusion criteria to verify measurement of gene-expression from whole blood using either microarray or RNA-Seq. The secondary filtering step included a detailed review of the metadata on Gemma and published methodology. Datasets that did not include a chronic stress manipulation were excluded. Abbreviations: n=number of datasets.

Three datasets met inclusion/exclusion criteria (GSE68076, GSE72262, GSE84185). **Table 1** summarizes the features of each experiment. Each study exposed eight-week old mice to chronic stress and profiled the whole blood transcriptome using Agilent microarray. Mice in GSE68076 were exposed to either 5 or 10 days of SDS and blood collection occurred either one, ten, or 42 days after last stress exposure ^11^. Mice in GSE72262 and GSE84185 experienced six weeks chronic mild stress, or eight weeks unpredictable chronic mild stress, respectively, with blood collection one day after final stress exposure.

**Table 1.** Overview of Studies Included in the Meta-Analysis. The sample size (“n”) only includes data from whole blood and excludes outlier samples. The Author and Year refer to the publication ^11–13^. All experimental details refer specifically to the samples used in the transcriptional profiling experiment.

| Project and Reference | Study Title | Subjects | Stress Exposure | Other Manipulated Variable | Time of Sacrifice | Array |
| --- | --- | --- | --- | --- | --- | --- |
| GSE68076<br>Muhie et.al.,<br>2017 | Molecular indicators of stress-induced neuroinflammation in a mouse model simulating features of post-traumatic stress disorder | Male C57Bl6<br>8–10-week-old<br>n = 25<br>Single Housed | Social Defeat Stress (SDS)<br>5 or 10 days | Stress duration and delay to sacrifice | 1 or 10days post 5-day SDS<br>1, 10, or 42 days post 10-day SDS | Agilent Genome-wide Mouse Expression arrays (GE_4x44Kv2 two-color) |
| GSE72262<br>Miyata et.al.,<br>2016 | Gene expression alterations in the medial prefrontal cortex and blood cells in a mouse model of depression during menopause | Female C57Bl6/J<br>8-week-old<br>n = 24<br>Housed 6 / cage | Chronic mild stress (CMS)<br>6 weeks | Bilateral ovariectomy or sham surgery 2 weeks prior to stress. | 1 day post final stress | Agilent SurePrint G3 Mouse GE 8 × 60 K Microarray (Design ID: 028005) |
| GSE84185<br>Herve et.al.,<br>2017 | Translational Identification of Transcriptional Signatures of Major Depression and Antidepressant Response | Male Balb/c<br>8-week-old<br>n = 32<br>Housed 4 / cage | Unpredictable chronic mild stress (UCMS)<br>8 weeks | Fluoxetine or control treatment during weeks 3-8 of UCMS; 10-20mg/kg/day | 1 day post final stress | Agilent Sure Print G3 Mouse Gene Expression 8x60K Microarray v1 |

### Individual Dataset Preprocessing

Probe alignment to an updated genome and manual curation for common issues was performed by Gemma^24,25^. Matrices containing annotated Log2 gene expression for each sample were accessed using *Gemma*.*R* (v.1.3.2). Each dataset was pre-processed to subset relevant samples (blood) and exclude genes lacking variation. Quality control included verifying Log2 transformation and identifying outliers using sample-sample correlations, following criteria adapted from Gemma ^24^. The final sample size was *n*=92 (*n*=45 Non-Stress/*n*=47 Stress: *GSE68076*: *n*=17 Non-Stress/n=19 Stress (1 outlier per condition eliminated), *GSE72262*: *n*=12 Non-Stress/n=12 Stress (no outliers), *GSE84185*: *n*=16 Non-Stress/*n*=16 Stress (no outliers). Potentially impactful technical variables (*e*.*g*., batch, library size), were screened for relationships with the top four principal components of variation in the expression data (function: *prcomp(x, scale=TRUE)*, package: *stats*). Differential expression was calculated using Limma (v.3.58.1 ^26^) with empirical Bayes moderation.

**Eq.1. Differential Expression Model for GSE68076**

*y = model*.*matrix(Stress_Factor + StressDuration_Factor10 + DelayToSample_Numeric* Animals from all conditions were included because all showed behavioral stress effects (reference levels: *Stress_Factor*: “non stress”, *StressDuration_Factor*: “5 days”).

**Eq.2 Differential expression model for GSE72262**

*y = model*.*matrix(Stress_factor+ OvX_factor)* Both intact and OvX animals were included to increase the generalizability of results (reference levels: *Stress_factor:* “non stress”, *OvX_factor*: “sham”).

**Eq.3 Differential expression model for GSE84185**

*y = model*.*matrix(~Stress_factor + Drug_factor + Stress_factor*Drug_factor)* All animals were included, with the modulation of stress effects by fluoxetine modeled as an interaction term to increase the power to capture stress effects in vehicle-treated animals and stress effects common to both treatment groups (reference levels: *Stress_factor:* “non stress”; *Drug_factor:* “vehicle”).

### Meta-Analysis

Results from the datasets were aligned by Entrez Gene ID. For genes with results from all three studies, a simple (intercept-only) random effects meta-analysis model was fit to the stress effect sizes (Log(2) Fold Change values or Log2FC) and their accompanying sampling variance (function *rma()*, package: *metafor* v.4.4-0 ^27^). False discovery rate (FDR) correction was performed (package: *multtest* v.2.58.0; method: Benjamini-Hochberg ^28–30^, with differentially expressed genes (DEGs) defined as FDR<0.05.

### Functional Ontology

Functional patterns were assessed with fast Gene Set Enrichment Analysis (*fgsea* v.1.28.0 ^31^; using a gene set database including brain-specific gene sets (Brain.GMT v.2) and traditional gene ontology (MSigDB: “C5: GO Gene Sets”) ^32–34^. fGSEA was performed using gene symbols ranked by the average estimated Log2FC.

### Follow-up Analysis: Exploring DEG Enrichment in Blood Cell Types

#### scRNA-Seq Dataset

To investigate which blood cell types express the identified stress DEGs, we used a single-cell RNA-Seq dataset (10xGenomics platform, iQ Biosciences) profiling peripheral blood mononuclear cells from C57BL/6 mice with a publicly-available raw Gene Expression library (Illumina NovaSeq 6000 ^35^).

#### scRNA-Seq Data Preprocessing

Analysis followed a Seurat pipeline ^36^ (adapted from: ^37^). Droplets were excluded with insufficient depth (<500 features), doublets/multiplets (>5,000 features), or poor health (>5% mitochondrial transcripts), leaving 4,058 high-quality cells. Gene expression was normalized by dividing each feature count by the total counts per cell, scaled by 10,000, and log-transformed (function: *NormalizeData()*).

To identify cell type clusters, dimensionality reduction (function: *RunPCA()*) was performed following Z-score normalization (function: *ScaleData()*) using data from 5,000 genes with the greatest standardized variance (function: *FindVariableFeatures()*). Clusters were visualized using the first 10 principal components and t-distributed Stochastic Neighbor Embedding (function: *RunTSNE()*) and Uniform Manifold Approximation (function: *RunUMAP())*. A Shared Nearest Neighbor graph was constructed (function: *FindNeighbors()*), followed by Louvain community detection (function: *FindClusters()*, resolution=1). Cluster-specific marker genes were identified using the Wilcoxon Rank Sum test (function: *FindAllMarkers()*, only.pos = TRUE, thresh.use=0.5, min.pct=0.25). Cell type identities were assigned using scCatch ^38^ (functions: *cellmatch()* and *findcelltype()*, parameters: species=“Mouse”, tissue=“Peripheral blood” or “Blood”), and validated and refined using published markers ^39–41^.

## Results

### Differentially Expressed Genes (DEGs)

There were 9,221 genes represented in all three studies, 9,219 of which produced stable meta-analysis estimates. Of these, 40 were differentially expressed (FDR<0.05) with 23 DEGs downregulated by stress (**Table 2a;** *Examples:* **Figure 2**) and 16 DEGs upregulated by stress (**Table 2b;** *Examples:* **Figure 3**). Forest plots illustrating the effects for all DEGs can be found in **Figure S1**. See **Table S1** for full statistical reporting (all 9,219 genes).

**Table 2.**
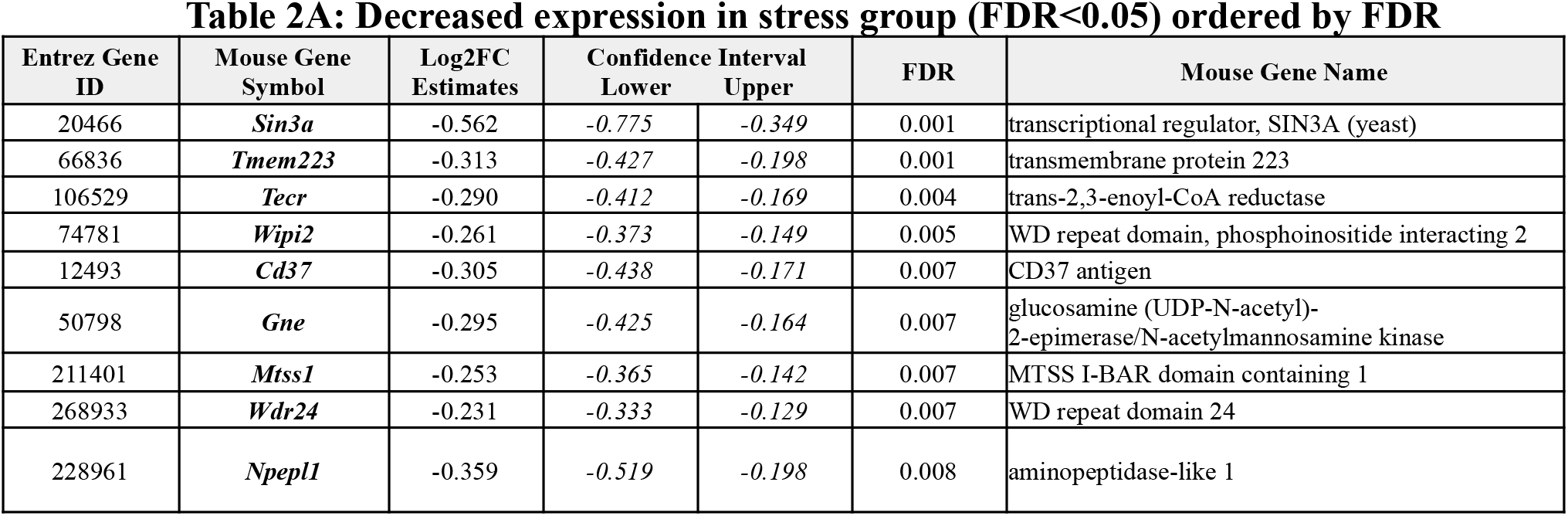
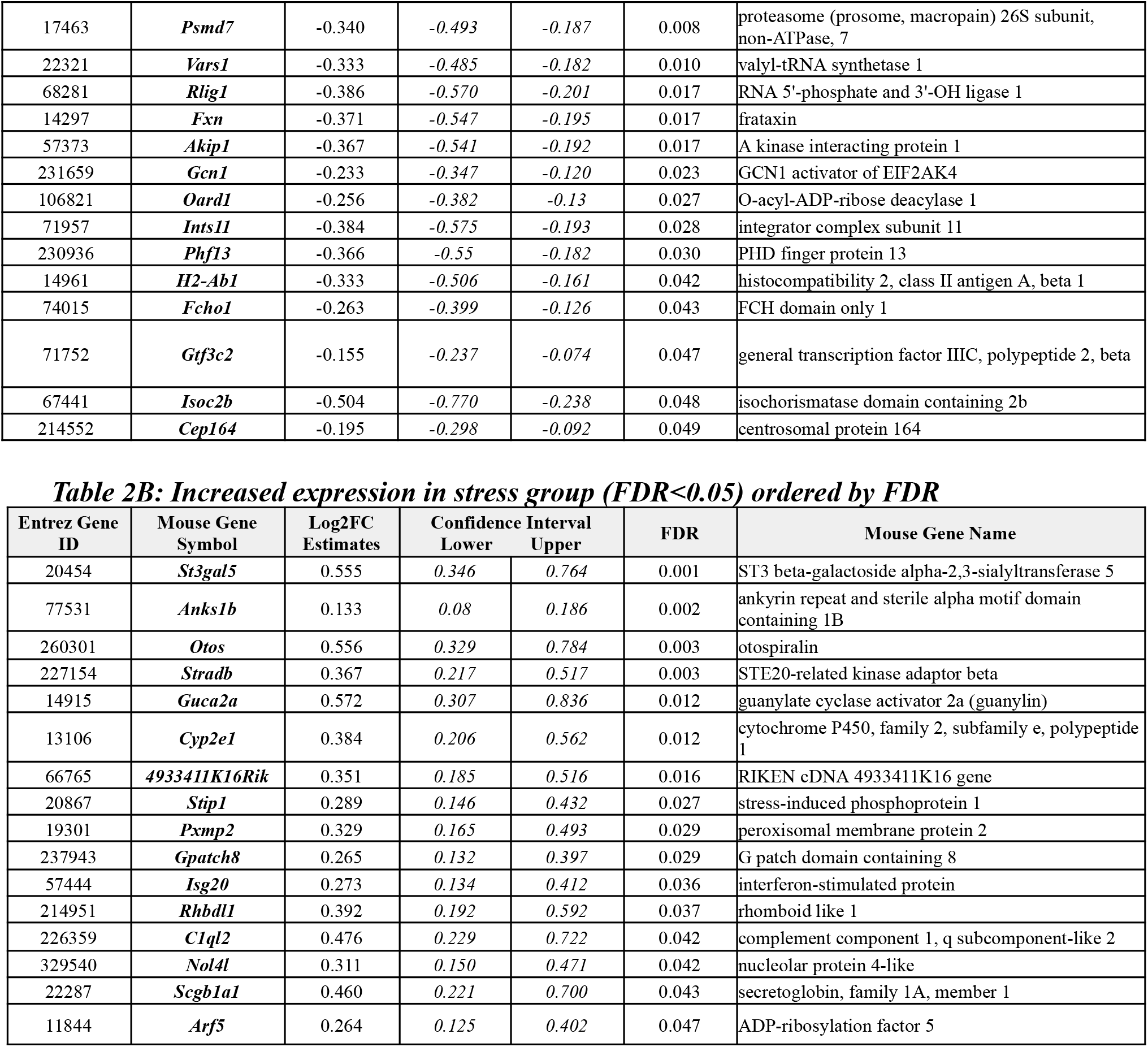
Meta-Analysis Results: Top Differentially Expressed Genes (DEGs: FDR<0.05) in whole blood of rodents exposed to chronic stress. The rows are ordered by FDR (lowest to highest). Definitions: Log2FC = Log2 Fold Change (Chronic Stress vs. Control), p-value = nominal p-value, FDR = False Discovery Rate (q-value). A negative Log2FC indicates lower expression in the experimental group relative to the control (downregulated; Table 2A), where larger negative values signify more substantial decreases. A positive Log2FC denotes that a gene’s expression is higher in the experimental group compared to the control group (upregulated; Table 2B), with higher positive values indicating greater increases.

**Figure 2.**
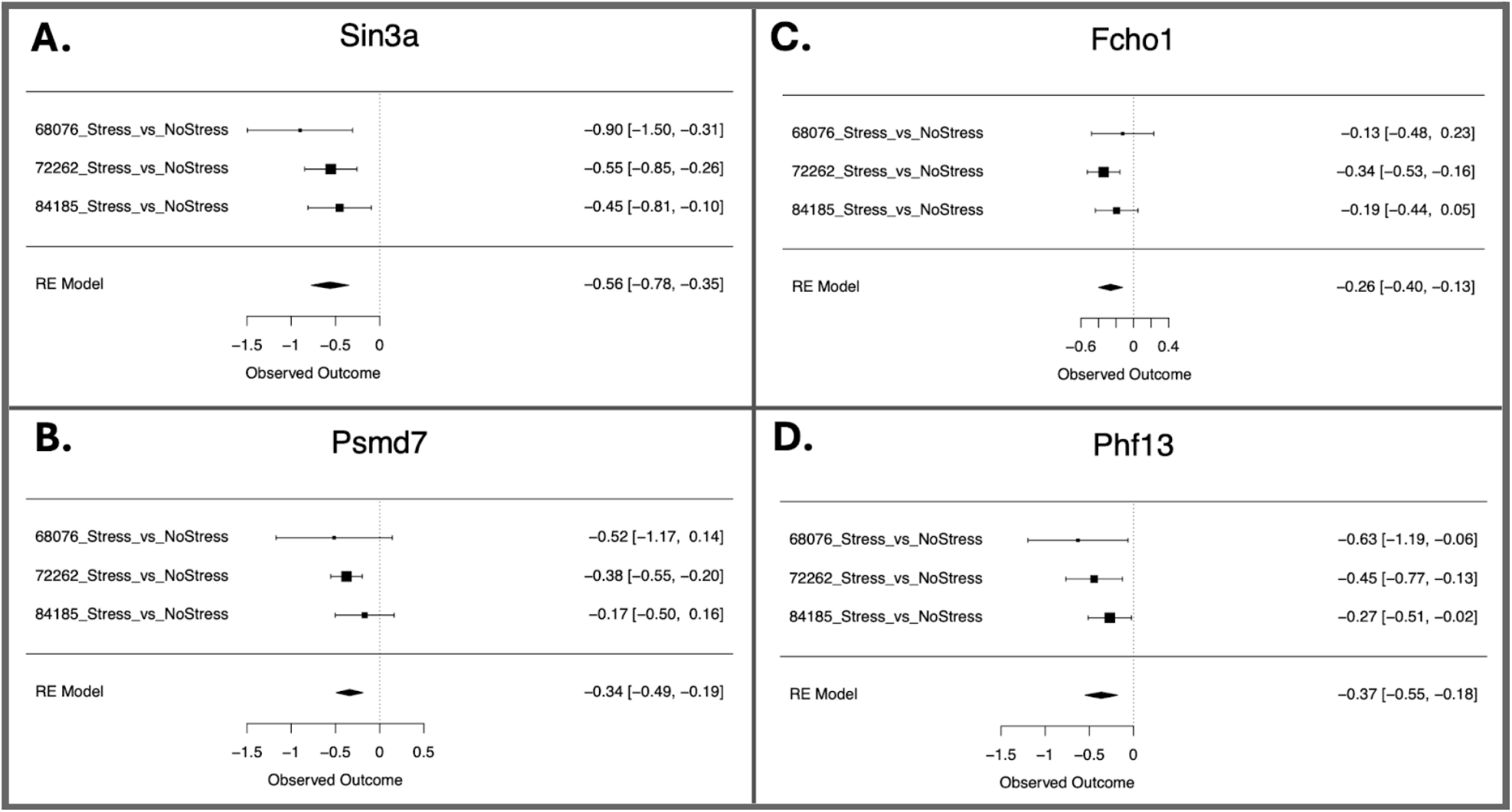
Multiple DEGs that were downregulated in response to chronic stress are associated with both immune function and processes related to chromatin organization and the cell cycle. A-D. Forest plots for differentially expressed genes that were down-regulated in response to chronic stress within the meta-analysis (FDR<0.05). Rows illustrate chronic stress Log2FC (squares) with 95% confidence intervals (whiskers) for each of the datasets and the meta-analysis random effects model (“RE Model”). Forest plots allow for visual inspection of the consistency and magnitude of effects across the three studies. Downregulated genes have a negative observed outcome for the RE model. **A**. Sin3a (transcriptional regulator, SIN3A (yeast)) **B**. Psmd7 (proteasome (prosome, macropain) 26S subunit, non-ATPase, 7) **C**. Fcho1 (FCH domain only 1) **D**. Phf13 (PHD finger protein 13)

**Figure 3.**
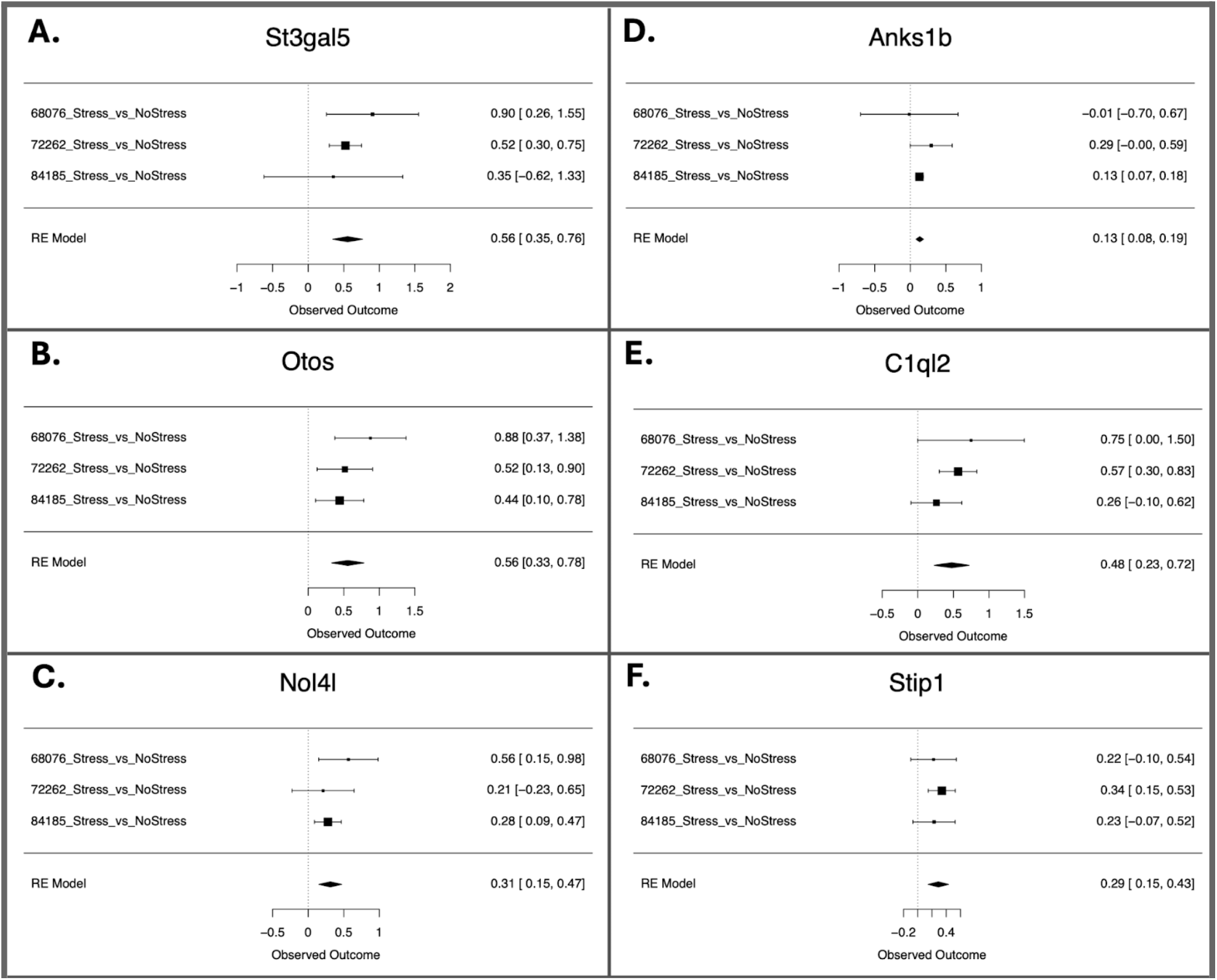
Many DEGs that were upregulated in response to chronic stress are associated with nervous system cell types, functions, and disorders. A-F. Forest plots for differentially expressed genes that were upregulated in response to chronic stress in the meta-analysis (FDR<0.05). Rows illustrate chronic stress Log2FC (squares) with 95% confidence intervals (whiskers) for each of the datasets and the meta-analysis random effects model (“RE Model”). Forest plots allow for visual inspection of the consistency and magnitude of effects across the three studies. Upregulated genes have a positive observed outcome for the RE model. **A**. St3gal5 (ST3 beta-galactoside alpha-2,3-sialyltransferase 5) **B**. Otos (otospiralin) **C**. Nol4l (nucleolar protein 4-like) **D**. Anks1b (ankyrin repeat and sterile alpha motif domain containing 1B) **E**. C1ql2 (complement component 1, q subcomponent-like 2) **F**. Stip1 (stress-induced phosphoprotein 1)

### Functional Gene Ontology

fGSEA output included 8,091 total gene sets. Of these, 212 were downregulated following stress and 60 gene sets were upregulated (FDR<0.05). Representative results are shown in **Table S2A&B**. See **Table S3** for the full statistical reporting (all 8,091 gene sets). In brief, DEGs downregulated in the meta-analysis are leading genes in enriched pathways related to DNA, chromosome organization, the cell cycle, gene silencing and epigenetic modifications (*Cep164, Oard1, Phf13, Psmd7, Sin3a*). Downregulated DEGs are also found in pathways related to gene-expression at the levels of mRNA production and processing (*Gcn1, Gtf3c2, Ints11, Psmd7, Sin3a*) and protein synthesis, metabolism, and modification (*Gcn1, Gne, Psmd7, Sin3a, Tecr*). Downregulated DEGs are involved in mitochondrial organization (*Fxn, Wipi2*), positive regulation of immune response, immunoglobulin production, adaptive immune response (*H2-Ab1, Psmd7, Sin3a*) and microglia (*Tecr*). Interestingly, several downregulated DEGs were implicated in abnormal development of midline structures and related functions (*Cep164, Fxn, Gne, H2-Ab1, Sin3a, Tecr*) and neurological problems linked to Alzheimer’s disease (*Cd37, Gtf3c2, Wipi2*), speech impairment (*Fxn, H2-Ab1, Tecr, Wipi2*), motor tone and tendon reflexes (*Gne, Sin3a, Tecr*; *Fxn, H2-Ab1*), and neuroblastoma (*Phf13, Wdr24*), suggesting broad relevance to neural development or function. DEGs upregulated following chronic stress were enriched in synapses (*Anks1b*), plasma membrane (*Rhbdl1, St3gal5*), and midbrain neurotypes (*Anks1b, C1ql2, Nol4l, St3gal5*). Upregulated DEGs were also implicated in oxygen binding (*Cyp2e1*), hormone activity (*Guca2a*), nervous system processes (*Otos*), and negative regulation of apoptotic signaling (*Stradb*).

### Exploring DEG enrichment in blood cell types

Within the fGSEA results, the gene set*”Bone marrow follicular B cell”* was enriched with stress-related down-regulation. As whole blood samples included many cell types, we decided to explore which blood cell types express each of the chronic stress DEGs using publicly available scRNA-Seq data from mouse peripheral blood mononuclear cells (PBMCs). PBMC samples contain most blood cell types, including lymphocytes (T cells, B cells, NK cells), monocytes, and dendritic cells. Amongst our DEGs, four had elevated relative expression in B cell clusters: *H2-Ab1* (*Log2FC*=1.15, *FDR*=1.36E-126, **Figure 4A**), *Cd37* (*Log2FC*=1.03, *FDR*=1.07E-63, **Figure 4B**), *Stip1* (*Log2FC*=0.51, *FDR*=8.50E-06), and *Oard1* (*Log2FC*=0.52, *FDR*=0.00020, **Figure 4C**). An additional DEG had highly elevated relative expression in an adjacent cluster characterized by both B cell and stem cell markers (*Mtss1: Log2FC*=3.05, *FDR*=3.45E-53, **Figure 4D**). The differential expression of these DEGs following chronic stress may particularly affect B Cell function.

**Figure 4.**
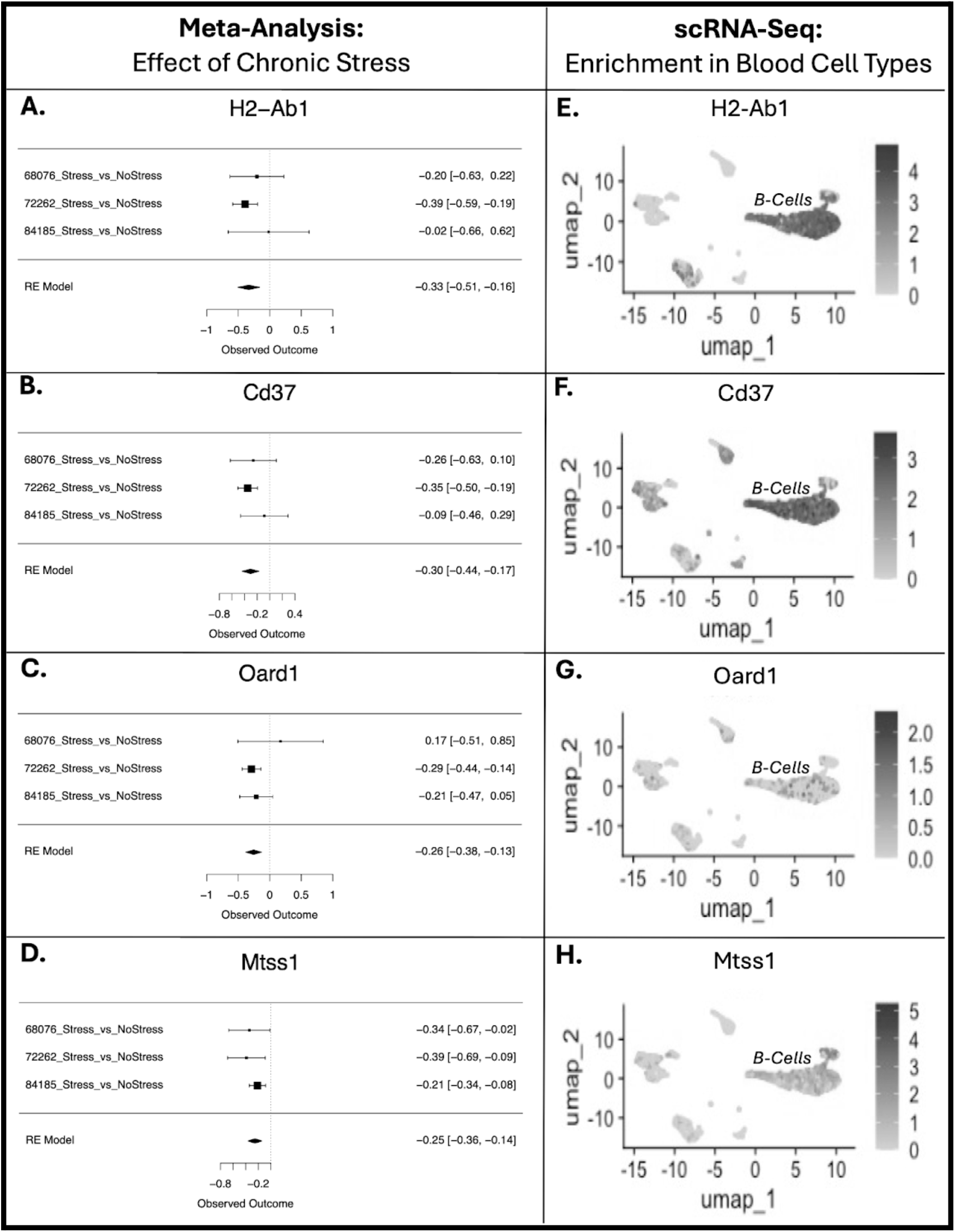
Multiple genes that were downregulated in response to chronic stress have highly enriched expression in B-Cells. A-D. Forest plots for differentially expressed genes that were down-regulated in response to chronic stress within the meta-analysis (FDR<0.05). Rows illustrate chronic stress Log2FC (squares) with 95% confidence intervals (whiskers) for each of the datasets and the meta-analysis random effects model (“RE Model”). Forest plots allow for visual inspection of the consistency and magnitude of effects across the three studies. Downregulated genes have a negative observed outcome for the RE model. **E-H**. Each of the DEGs illustrated in A-D have elevated expression in B Cells under basal conditions within a publicly available scRNA-Seq dataset from mouse peripheral blood mononuclear cells (PBMCs). Shown are UMAP plots illustrating the cell type clusters (x-axis: UMAP dimension 1, y-axis: UMAP dimension 2), with shading intensity (grey) indicating Log expression for the gene of interest. **A**. H2-Ab1 (histocompatibility 2, class II antigen A, beta 1) **B**. Cd37 (CD37 antigen) **C**. Oard1 (O-acyl-ADP-ribose deacylase 1) **D**. Mtss1 (MTSS I-BAR domain containing 1).

## Discussion

Animal models in stress neurobiology research promise to improve diagnosis and treatment for humans. To date, this promise has not fully reached fruition. As blood samples are accessible in both animal models and humans with stress-sensitive disorders, examining the effects of chronic stress on the blood transcriptome can provide an important cross-species bridge. By statistically combining three smaller studies with different chronic stress paradigms, our study improves the generalizability of our findings illustrating the effects of chronic stress on blood, while also increasing statistical power and minimizing the false positive risk. Our meta-analysis revealed that transcripts down-regulated following chronic stress were related to B cells, immune function, DNA/chromatin regulation, RNA processing and translation, cellular metabolic processes, and mitochondrial function; upregulated transcripts related to erythrocytes and oxygen binding, synapses, cell junctions, transport, and cell signaling. Potential implications of these findings are discussed below.

### Inflammation and Immunodeficiency

Chronic stress is well known to contribute to inflammation and disrupt innate and adaptive immunity ^42,43^. Several of the stress-responsive DEGs identified in our meta-analysis may contribute to these effects. For example, gene sets associated with B lymphocytes (B cells) were downregulated in response to chronic stress. B cells are important for the production of antibodies, immune memory, antigen presentation, and secretion of cytokines and trophic factors. B cells also possess neurotransmitter receptors, allowing for bidirectional communication with the nervous system. Stress is known to modify B cell development and survival, cell-cell interaction, and migration into brain or brain-adjacent tissue ^44^. Several stress-responsive DEGs in our meta-analysis had highly enriched expression in B-cells within blood scRNA-Seq data, including *H2-Ab1, Cd37, Stip1*, and *Oard1*. Differential expression of these genes following stress may influence B cell function. For example, *Cd37* influences B-cell/T-cell interactions, balances immunoglobulin (Ig) IgG versus IgA production, and limits interleukin (IL-2, IL-4) production and T-cell proliferation. Decreased *Cd37* expression following stress could increase susceptibility to pathogens, especially given the importance of IgG in blood ^45^. *Mtss1* was down-regulated following chronic stress, and highly expressed in immature B cells in blood scRNA-Seq data. Decreased *Mtss1* can disrupt B-cell receptor signaling by decreasing IgG2b and compromise IgM response to antigens ^46^.

Inflammation may follow other stress-responsive differential expression. Several gene sets related to immune system regulation were down-regulated following stress, including genes *Sin3a, Psmd7*, and *H2-Ab1*. Decreased *Sin3a* could cause immune dysregulation as it helps maintain tight control over innate and adaptive immune factors ^47,48^. Likewise, decreased *Fcho1* can produce T-cell deficiency and immunodeficiency ^49^. Decreased *Phf13* may weaken immune response, as its protein (SPOC1) has anti-adenovirus properties ^50^. Stress-related upregulation of *Isg20* may distort innate immune response by B-and T-cells ^51^. Altogether, our results provide compelling candidates for mediating the effects of chronic stress on the immune system.

### Stress-Induced Cellular Aging

The meta-analysis results are consistent with previous reports linking chronic stress to accelerated cellular aging characterized by DNA damage, failed DNA repair, altered chromatin structure and cell cycle, disrupted protein homeostasis and quality control, and mitochondrial dysfunction ^52,53^.

Chronic stress exposure may increase DNA damage and hinder DNA repair secondary to altered signaling by stress hormones: cortisol, epinephrine, and norepinephrine ^54^. Several of the identified DEGs may mediate this relationship. Gene sets related to the cell cycle and G2M phase transition were down-regulated following stress, including genes *Fcho1, Sin3a, Psmd7*. Of particular relevance to blood, *Ints11* was also down-regulated, and regulates gene-expression for hematopoiesis; with genetic deficiency driving lethal arrest ^55^. Gene sets related to chromatin organization were also down-regulated, including DEGs important for early signalling of DNA damage (*Cep164* ^56^), DNA damage response (*Oard1* ^57^), and modulation of DNA damage response elements (*Phf13* ^58^). Many of these DEGs are also involved in immune response, since viral infection can necessitate DNA repair ^50^.

Our meta-analysis results support the hypothesis that chronic stress accelerates age-related decline of protein homeostasis and proteome quality control ^59,60^. Decreased *Gcn1* following stress could decrease activation of the integrated stress response, loosening cellular control on protein production ^61^. Stress induction of *Stip1* can disrupt protein folding and proteolytic degradation ^62^. Additional disruptions in protein removal and recycling may follow downregulation of DEGs encoding deubiquitinase PSMD7 and aminopeptidase NPEPL1. Failure to eliminate problematic proteins may be exacerbated by decreased *Wdr24*, which mediates protein homeostatic processes including detection of amino acid starvation, ubiquitin protein ligase activity, and lysosomal acidification ^63^. Stress effects on protein homeostasis may be exacerbated by decreased *Vars1*, which encodes valyl-tRNA synthetase ^64^.

Cellular aging could be further promoted by stress-induced mitochondrial dysregulation following down-regulation of *Tmem222, Akip1, Fxn, and Wipi2. Tmem223* supports mitochondrial cytochrome c oxidase assembly ^65^, and *Akip1* is critical for cellular respiration and ATP production in some tissues ^66^. Decreased *Wipi2* and *Fxn* could destabilize mitochondrial copy number, due to involvement in mitophagy ^67,68^ and mitochondrial biogenesis ^69^. Overall, these effects of chronic stress on processes related to cellular aging may, in part, explain the wide range of symptoms and disorders prevalent in people with a history of chronic stress or trauma.

### Links to Brain-Based Disorders

Several of the stress-sensitive DEGs were previously implicated in brain-based disorders, suggesting potential relevance to stress-induced psychopathology. For example, stress-induced downregulation of *Tmem223* resembles the downregulation seen in blood mononucleocytes in a rodent model of schizophrenia ^70^, suggesting utility as a psychiatric biomarker. Our work confirms prior findings showing stress-induced down-regulation of *Tecr*, a leading gene in pathways related to microglia and abnormal development. Decreased *Tecr* has been implicated in blood brain barrier degradation ^71^, with mutations causing non-syndromal mental retardation ^72^.

Gene sets related to synapses and midbrain neurotypes were enriched with stress-related differential expression. These findings may mirror central effects, as brain-based transcripts can be measured in blood due to exosomes ^5^. A leading DEG was *Anks1b*, which was upregulated following chronic stress. ANKS1B is ubiquitous in the brain as a postsynaptic scaffold protein at excitatory synapses ^73^ and implicated in disordered social behavior ^74^. ANKS1B may influence corticosteroid response ^73^, positioning it to mediate stress effects on excitatory neurotransmission.

Stress-induced disruption of protein homeostasis may also increase the risk for neurodegenerative disorders. Elevated *Stip1* - as observed following chronic stress - is linked to aggregation of misfolded proteins in animal models ^62^ and neurodegenerative disorders ^75^. Together these observations support a framework for embodied symptoms of stress-sensitive disorders.

### Strengths and Limitations

By performing a meta-analysis to statistically combine results from smaller studies using different chronic stress paradigms, we improve the generalizability of our findings, while increasing sample size and reducing false positive risk. Therefore, it is worth noting where our findings both converge and diverge from the conclusions of the individual studies included in the meta-analysis. In general, many pathways highlighted in our results were previously identified in one of the included studies, including mitochondrial function ^11–13^, immune function ^11,12^, protein ubiquitination ^12,13^, oxidative phosphorylation ^13^, RNA processing ^12^, and stress-induced cellular aging ^11^. Despite similarities in identified pathways, there was little overlap in specific DEGs (0.4% overlap with ^13^; 0% overlap with ^12^ ^11^).

These differences highlight the importance of meta-analyses as an antidote to the replication crisis by improving the power to detect smaller, consistent effects across studies and paradigms, reducing false positives and expanding external validity. That said, the power and generalizability of our findings are still limited because only three studies survived *a priori* inclusion/exclusion criteria and quality standards. As relevant future datasets are released, updating our meta-analysis may result in additional findings. Likewise, we were able to provide additional cell type information by cross-referencing our findings with public scRNA-Seq data from basal subjects, but inferences would be improved by direct measurements of stress effects on individual blood cell types using cell sorting or single cell sequencing.

## Conclusion

Chronic stress increases the risk for numerous somatic and psychiatric disorders. In addition to direct impacts on individuals, stress-sensitive disorders cost society lost productivity and medical expenses. Translation from animal models to human diagnostics and treatments has been slow and included high profile failures (*e*.*g*., CRF receptor antagonism ^76^. Our meta-analysis supports and extends previous findings by identifying converging effects across chronic stress paradigms and increasing statistical power to detect more moderate effect sizes. Our meta-analysis results show that chronic stress disrupts gene expression contributing to immune regulation, DNA stability, gene-expression, and protein homeostasis and supports the hypothesis that chronic stress accelerates cellular aging, which can contribute to brain inflammation and cognitive decline. That these pathways are relevant across neuropsychiatric diagnoses highlights the importance of interdisciplinary biomedical research. Our results should encourage future research toward accessible tissues like blood to directly compare metrics in human subjects and animal models and explore cellular mechanisms linking chronic stress to neuropsychiatric symptoms.

## Acknowledgements

This work was supported by the Hope for Depression Research Foundation (HDRF: HA), Grinnell College Center for Careers, Life, and Service (TQD, AB, & DMN), the International Brain Research Organization (IBRO) and Faculty for Undergraduate Neuroscience (FUN) (TQD & DMN), NIDA (U01 DA043098: HA), the Pritzker Neuropsychiatric Disorders Research Foundation (HA & SJW). Funders and sponsors had no active role in the review.

## CRediT statement

EIF: Conceptualization, Methodology, Software, Formal analysis, Investigation, Data Curation, Writing - Original Draft, Writing - Review & Editing, Visualization, Supervision

DMN: Conceptualization, Methodology, Software, Writing - Original Draft, Writing - Review & Editing

MHH: Conceptualization, Methodology, Software, Writing - Review & Editing, Visualization, Supervision, Project administration

MN: Methodology, Software, Writing - Review & Editing

HK: Methodology, Software, Writing - Review & Editing

TQD: Methodology, Software, Writing - Review & Editing

AB: Methodology, Software, Writing - Review & Editing

SJW: Writing - Review & Editing, Supervision, Funding acquisition

HA: Writing - Review & Editing, Supervision, Funding acquisition

