## Supplement for "Effect of Chronic Stress on Whole Blood Transcriptome: A Meta-Analysis of Publicly Available Datasets from Rodent Models"

### Supplemental Tables & Table Legends

***Table S1. The full meta-analysis results (9,221 genes, 9,219 stable meta-analysis estimates).*** *This .xlsx file includes two worksheets: 1) The worksheet "MetaAnalysisOutputByPval" provides the full meta-analysis results, with each row representing the results for one gene, and each column providing either gene annotation or meta-analysis statistical output. The results are ordered by p-value, so that the top rows in the worksheet are the genes with the smallest p-values. 2) The worksheet "ColumnDefinitions" provides the definitions for the variables present in each column in "MetaAnalysisOutputByPval"*

***Table S2A: Significantly enriched pathways; leading genes include those with decreased expression in mice exposed to chronic stress***

| Entrez ID | Gene Symbol | Representative Significantly Enriched Pathways (Chronic Stress) |
|---|---|---|
| 12493 | *Cd37* | HAY: Bone marrow follicular B cell |
| | | BLALOCK: Alzheimer's Disease Up |
| 214552 | *Cep164* | GOBP: DNA metabolic process, DNA repair |
| | | HP: Renal insufficiency, Hepatic Failure, Abnormal hepatobiliary system physiology |
| 74015 | *Fcho1* | GOBP: protein containing complex subunit organization |
| 14297 | *Fxn* | GOBP: Organonitrogen biosynthesis, Mitochondrion organization |
| | | GOCC: Mitochondrion, Mitochondrial Matrix |
| | | HP: Hypertrophic cardiomyopathy, abnormal myocardium morphology |
| | | HP: Abnormal Cellular Phenotype |
| | | HP: Hypertonia, Reduced Tendon Reflexes, Abnormal involuntary eye movements, Neurological speech impairment |
| 231659 | *Gcn1* | GOBP: Biosynthetic Process; Organonitrogen, Peptide, Cellular Amide |
| | | GOBP: Posttranscriptional regulation of gene expression |
| | | GOCC: Ribosome |
| | | GOMF: Ribonucleoprotein complex binding, RNA binding |
| 50798 | *Gne* | GOBP: Cellular amide metabolic process |
| | | HP: Reduced tendon reflexes, generalized hypotonia |
| | | HP: Abnormal Cellular Phenotype |
| | | HP: Abnormal myocardium morphology, elevated hepatic transaminase, abnormality of the liver |
| 71752 | *Gtf3c2* | GOBP: ncRNA Transcription |
| | | BLALOCK: Alzheimer's disease up |
| 14961 | *H2-Ab1* | GOBP: Adaptive immune response, Immunoglobulin production, lymphocyte mediated immunity, positive regulation of immune response |
| | | GOCC: Golgi apparatus subcompartment, Organelle subcompartment |
| | | HP: Hypertonia |
| | | HP: Unusual infection |
| 71957 | *Ints11* | GOBP: ncRNA metabolic process, ncRNA transcription, RNA processing, snRNA transcription |
| | | GOCC: Nuclear protein containing complex |
| 211401 | *Mtss1* | GOBP: endomembrane system organization |
| 228961 | *Npepl1* | GOMF: Manganese ion binding |
| 106821 | *Oard1* | GOBP: Cellular response to DNA damage, DNA conformational change, DNA packaging |
| | | GOCC: Nucleolus, Chromosome |
| | | GOMF: Methylated histone binding |

| | | |
|---|---|---|
| | | JOHNSTONE: ParvB targets 2 down |
| 230936 | *Phf13* | GOBP: Cell cyle, chromatin organization |
| | | GOMF: Chromatin binding |
| 17463 | *Psmd7* | GOBP: Cell cycle, G2M phase transition, macromolecule catabolic process, mRNA metabolic process, Negative regulation of gene expression, Post-transcriptional regulation of gene expression, RNA catabolic process |
| | | GOBP: Positive regulation of immune response, regulation of innate immune response |
| | | GOBP: Protein modification by small protein conjugation |
| | | GOCC: Catalytic complex, Intracellular protein containing complex |
| 20466 | *Sin3a* | GOBP: Cell cycle, G2M phase transition, cellular macromolecule localization, negative regulation of chromatin organization, gene silencing, protein localization to organelle, regulation of gene expression epigenetic, peptidyl lysine modification |
| | | GOBP: Positive regulation of immune response, regulation of immune effector process, regulation of innate immune response, |
| | | GOCC: Catalytic Complex, Chromosome, Chromosome centromeric region, kinetochore, nuclear protein containing complex |
| | | GOMF: Chromatin binding, RNA binding |
| | | HP: Generalized hypotonia, Abnormal development, feeding difficulties, facial shape, eye movements, head circumference |
| | | HP: Unusual infection |
| 106529 | *Tecr* | GOBP: Amide biosynthetic / metabolic process, organonitrogen biosynthetic process |
| | | GOCC: Organelle subcompartment |
| | | HP: Abnormal development of face and head; generalized hypotonia, neurological speech impairment |
| | | JOHNSTONE: ParvB Targets 2 down |
| | | ZHONG: PFC C1 microglia |
| 268933 | *Wdr24* | Park_2011_Coexpression_Hippocampus_Mouse_darkgrey |
| 74781 | *Wipi2* | BLALOCK: Alzheimer's disease up |
| | | GOBP: Cellular macromolecule localization, mitochondrion organization, organonitrogen compound biosynthetic process |
| | | HP: neurological speech impairment |

***Table S2B: Significantly enriched pathways; leading genes include those with increased expression in mice exposed to chronic stress***

| Entrez ID | Gene Symbol | Representative Enriched Pathways (Chronic Stress) |
|---|---|---|
| 77531 | *Anks1b* | GOCC: Synapse |
| | | MANNO: Midbrain neurotypes HDA1 |
| 226359 | *C1ql2* | MANNO: Midbrain neurotypes HGABA |
| | | MEISSNER: Brain HCP with H3K27ME3 |
| 13106 | *Cyp2e1* | GOMF: Oxygen binding |
| 14915 | *Guca2a* | GOMF: Hormone activity |

| | | |
|---|---|---|
| 329540 | *Nol4l* | MANNO: Midbrain neurotypes HDA1 |
| 260301 | Otos | GOBP: Nervous system process |
| 214951 | Rhbdl1 | GOCC: intrinsic component of plasma membrane |
| 20454 | St3gal5 | GOCC: intrinsic component of plasma membrane |
| | | MANNO: Midbrain neurotypes HDA1 |

***Table S2. Functional gene sets that are enriched with the effects of chronic stress.***
*fGSEA identified 212 gene-set pathways significantly downregulated in stress-exposed mice and 60 significantly upregulated gene-set pathways (FDR < 0.05). Table S2 displays significantly enriched biological pathways, molecular functions, and cellular components with leading genes that were significantly down- (Table S2A) or up-regulated (Table S2B) in our meta-analysis. Acronyms: GO: gene ontology, CC: cellular component, BP: biological process, MF: molecular function, HP: human phenotype*

***Table S3: The full fast Gene Set Enrichment Analysis (fGSEA) results (8,091 gene sets).*** *This .xlsx file includes two worksheets: 1) The worksheet "fGSEA_Results" provides the full fGSEA results, with each row representing the results for one gene set, and each column providing the fGSEA statistical output. The results are ordered by p-value, so that the top rows in the worksheet are the gene sets with the smallest p-values. 2) The worksheet "ColumnDefinitions" provides the definitions for the variables present in each column in "fGSEA_Results".*

### Supplementary Figures & Figure Legends

***Figure S1A: Forest plots of genes with decreased expression in mice exposed to chronic stress***

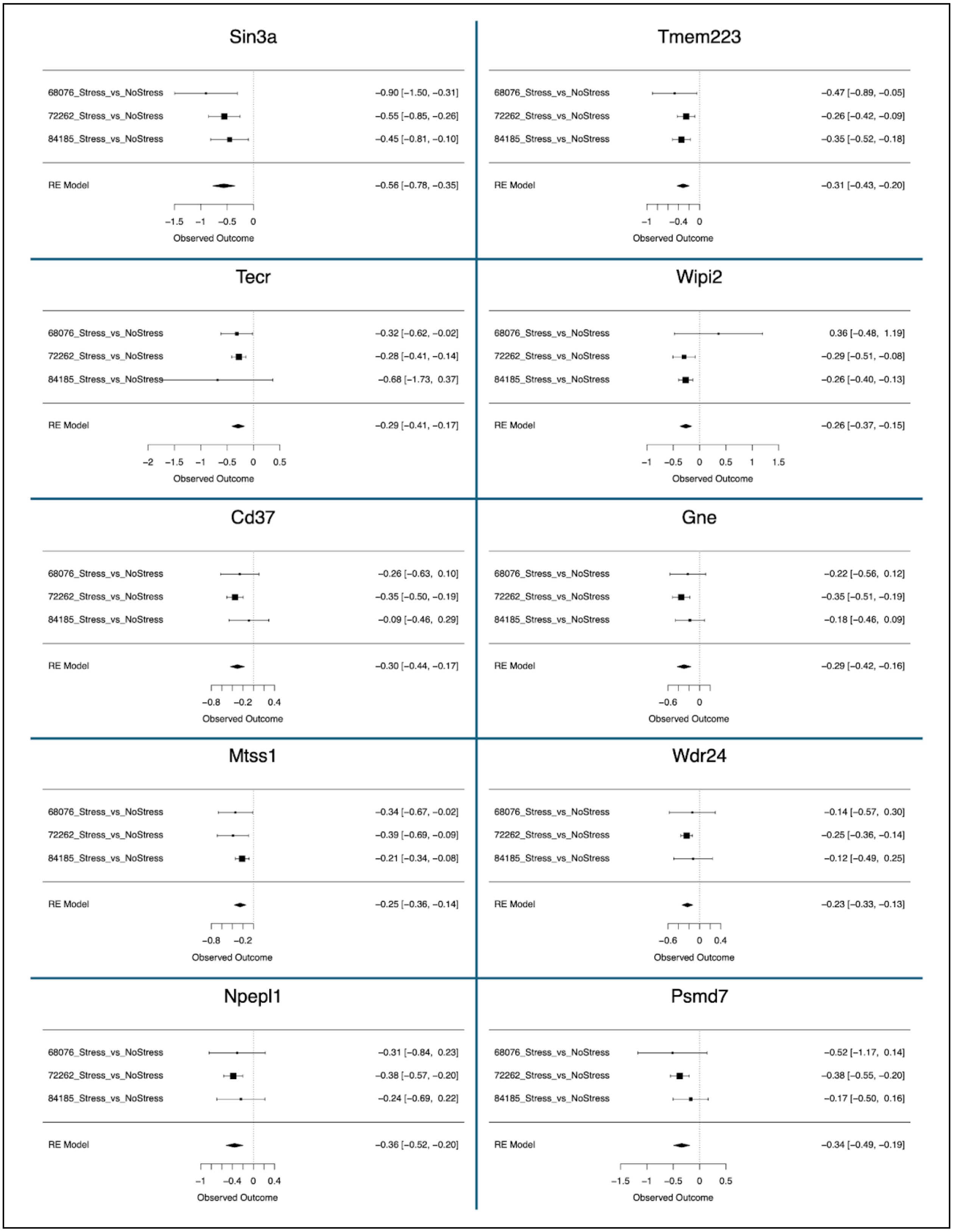

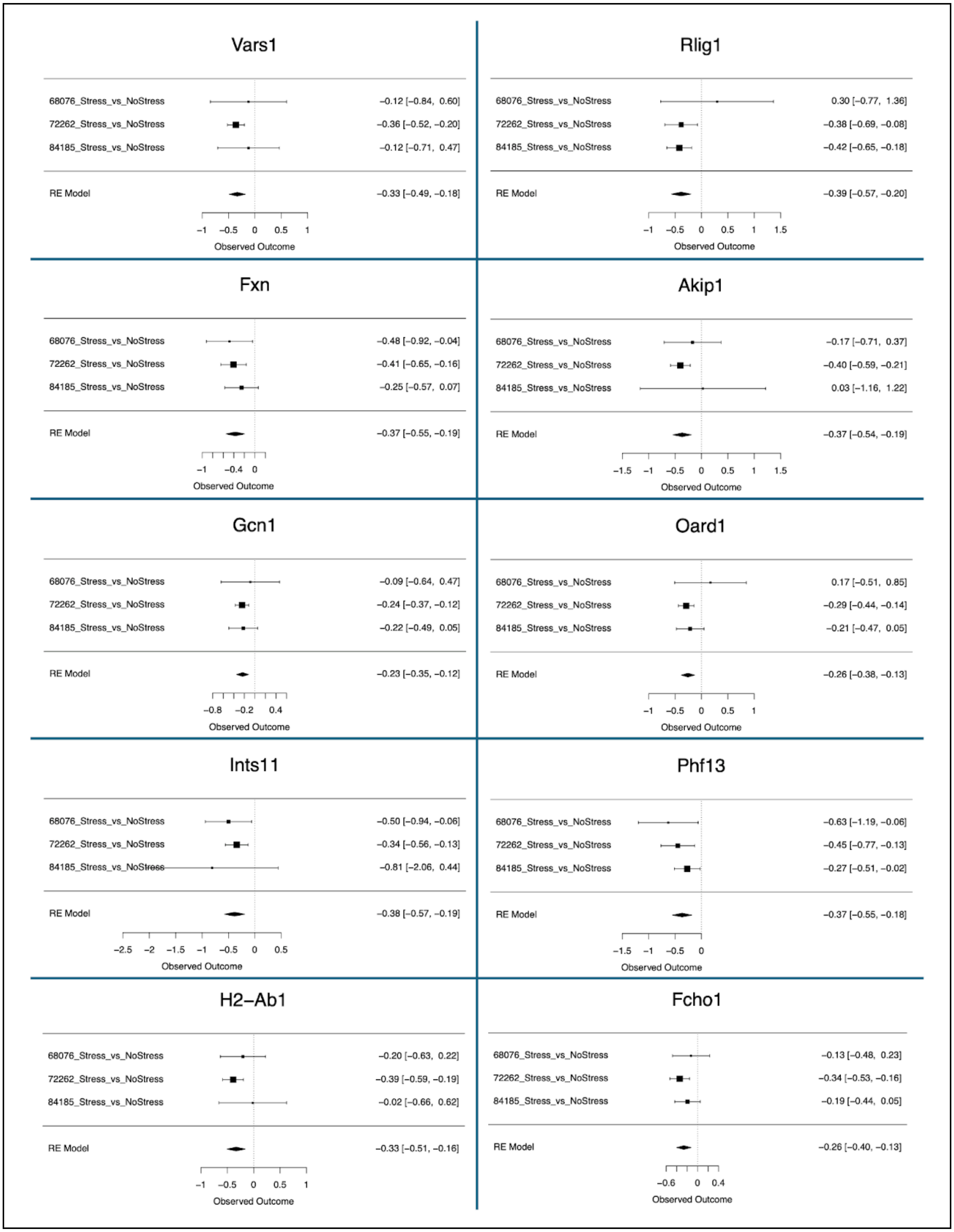
Vars1
68076_Stress_vs_NoStress −0.12 [−0.84, 0.60]
72262_Stress_vs_NoStress −0.36 [−0.52, −0.20]
84185_Stress_vs_NoStress −0.12 [−0.71, 0.47]
RE Model −0.33 [−0.49, −0.18]
Observed Outcome
Rlig1
68076_Stress_vs_NoStress 0.30 [−0.77, 1.36]
72262_Stress_vs_NoStress −0.38 [−0.69, −0.08]
84185_Stress_vs_NoStress −0.42 [−0.65, −0.18]
RE Model −0.39 [−0.57, −0.20]
Observed Outcome
Fxn
68076_Stress_vs_NoStress −0.48 [−0.92, −0.04]
72262_Stress_vs_NoStress −0.41 [−0.65, −0.16]
84185_Stress_vs_NoStress −0.25 [−0.57, 0.07]
RE Model −0.37 [−0.55, −0.19]
Observed Outcome
Akip1
68076_Stress_vs_NoStress −0.17 [−0.71, 0.37]
72262_Stress_vs_NoStress −0.40 [−0.59, −0.21]
84185_Stress_vs_NoStress 0.03 [−1.16, 1.22]
RE Model −0.37 [−0.54, −0.19]
Observed Outcome
Gcn1
68076_Stress_vs_NoStress −0.09 [−0.64, 0.47]
72262_Stress_vs_NoStress −0.24 [−0.37, −0.12]
84185_Stress_vs_NoStress −0.22 [−0.49, 0.05]
RE Model −0.23 [−0.35, −0.12]
Observed Outcome
Oard1
68076_Stress_vs_NoStress 0.17 [−0.51, 0.85]
72262_Stress_vs_NoStress −0.29 [−0.44, −0.14]
84185_Stress_vs_NoStress −0.21 [−0.47, 0.05]
RE Model −0.26 [−0.38, −0.13]
Observed Outcome
Ints11
68076_Stress_vs_NoStress −0.50 [−0.94, −0.06]
72262_Stress_vs_NoStress −0.34 [−0.56, −0.13]
84185_Stress_vs_NoStress −0.81 [−2.06, 0.44]
RE Model −0.38 [−0.57, −0.19]
Observed Outcome
Phf13
68076_Stress_vs_NoStress −0.63 [−1.19, −0.06]
72262_Stress_vs_NoStress −0.45 [−0.77, −0.13]
84185_Stress_vs_NoStress −0.27 [−0.51, −0.02]
RE Model −0.37 [−0.55, −0.18]
Observed Outcome
H2−Ab1
68076_Stress_vs_NoStress −0.20 [−0.63, 0.22]
72262_Stress_vs_NoStress −0.39 [−0.59, −0.19]
84185_Stress_vs_NoStress −0.02 [−0.66, 0.62]
RE Model −0.33 [−0.51, −0.16]
Observed Outcome
Fcho1
68076_Stress_vs_NoStress −0.13 [−0.48, 0.23]
72262_Stress_vs_NoStress −0.34 [−0.53, −0.16]
84185_Stress_vs_NoStress −0.19 [−0.44, 0.05]
RE Model −0.26 [−0.40, −0.13]
Observed Outcome

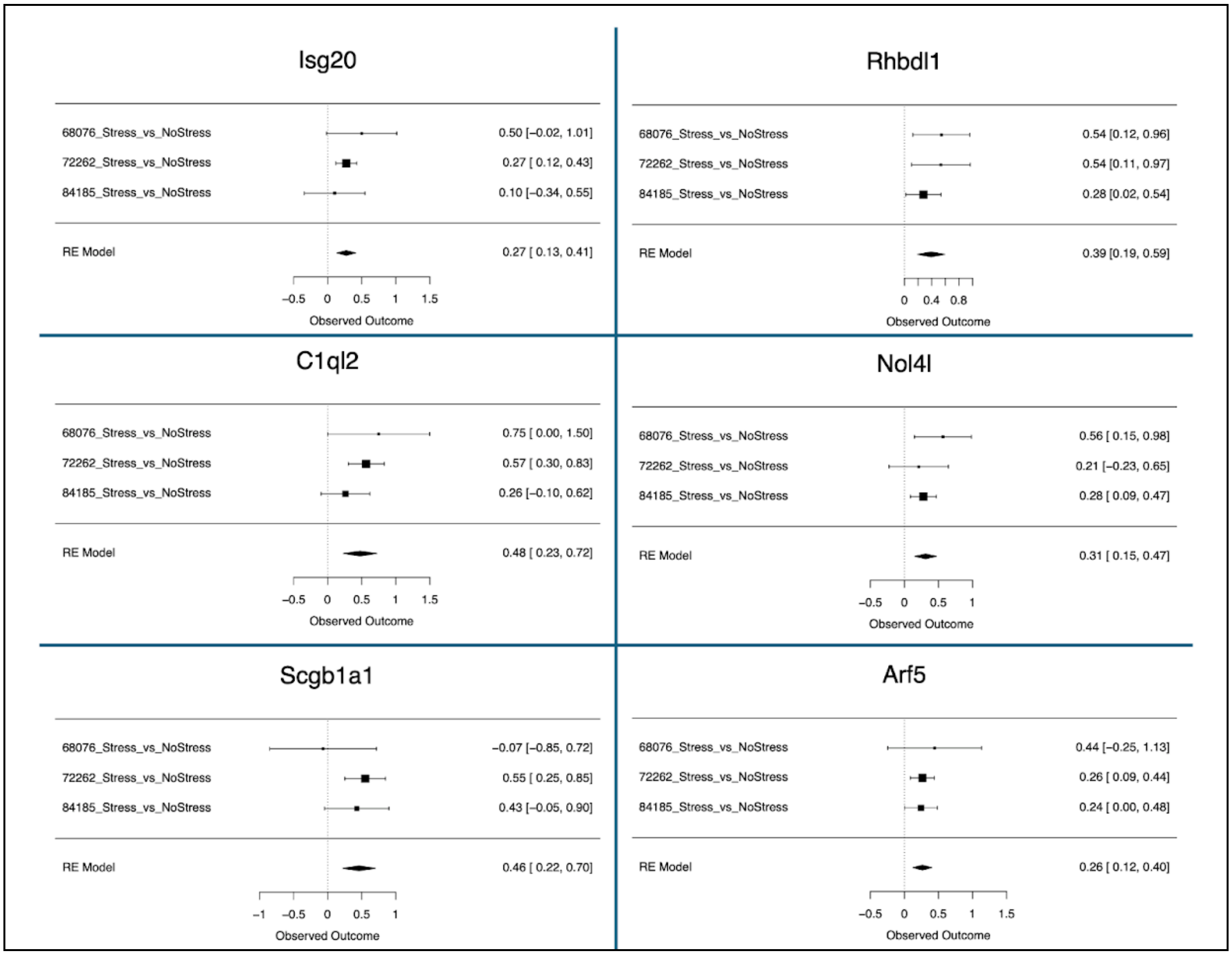

Isg20
68076_Stress_vs_NoStress 0.50 [−0.02, 1.01]
72262_Stress_vs_NoStress 0.27 [ 0.12, 0.43]
84185_Stress_vs_NoStress 0.10 [−0.34, 0.55]
RE Model 0.27 [ 0.13, 0.41]
Observed Outcome
Rhbdl1
68076_Stress_vs_NoStress 0.54 [0.12, 0.96]
72262_Stress_vs_NoStress 0.54 [0.11, 0.97]
84185_Stress_vs_NoStress 0.28 [0.02, 0.54]
RE Model 0.39 [0.19, 0.59]
Observed Outcome
C1ql2
68076_Stress_vs_NoStress 0.75 [ 0.00, 1.50]
72262_Stress_vs_NoStress 0.57 [ 0.30, 0.83]
84185_Stress_vs_NoStress 0.26 [−0.10, 0.62]
RE Model 0.48 [ 0.23, 0.72]
Observed Outcome
Nol4l
68076_Stress_vs_NoStress 0.56 [ 0.15, 0.98]
72262_Stress_vs_NoStress 0.21 [−0.23, 0.65]
84185_Stress_vs_NoStress 0.28 [ 0.09, 0.47]
RE Model 0.31 [ 0.15, 0.47]
Observed Outcome
Scgb1a1
68076_Stress_vs_NoStress −0.07 [−0.85, 0.72]
72262_Stress_vs_NoStress 0.55 [ 0.25, 0.85]
84185_Stress_vs_NoStress 0.43 [−0.05, 0.90]
RE Model 0.46 [ 0.22, 0.70]
Observed Outcome
Arf5
68076_Stress_vs_NoStress 0.44 [−0.25, 1.13]
72262_Stress_vs_NoStress 0.26 [ 0.09, 0.44]
84185_Stress_vs_NoStress 0.24 [ 0.00, 0.48]
RE Model 0.26 [ 0.12, 0.40]
Observed Outcome

***Figure S1B: Forest plots of genes with increased expression in mice exposed to chronic stress.***

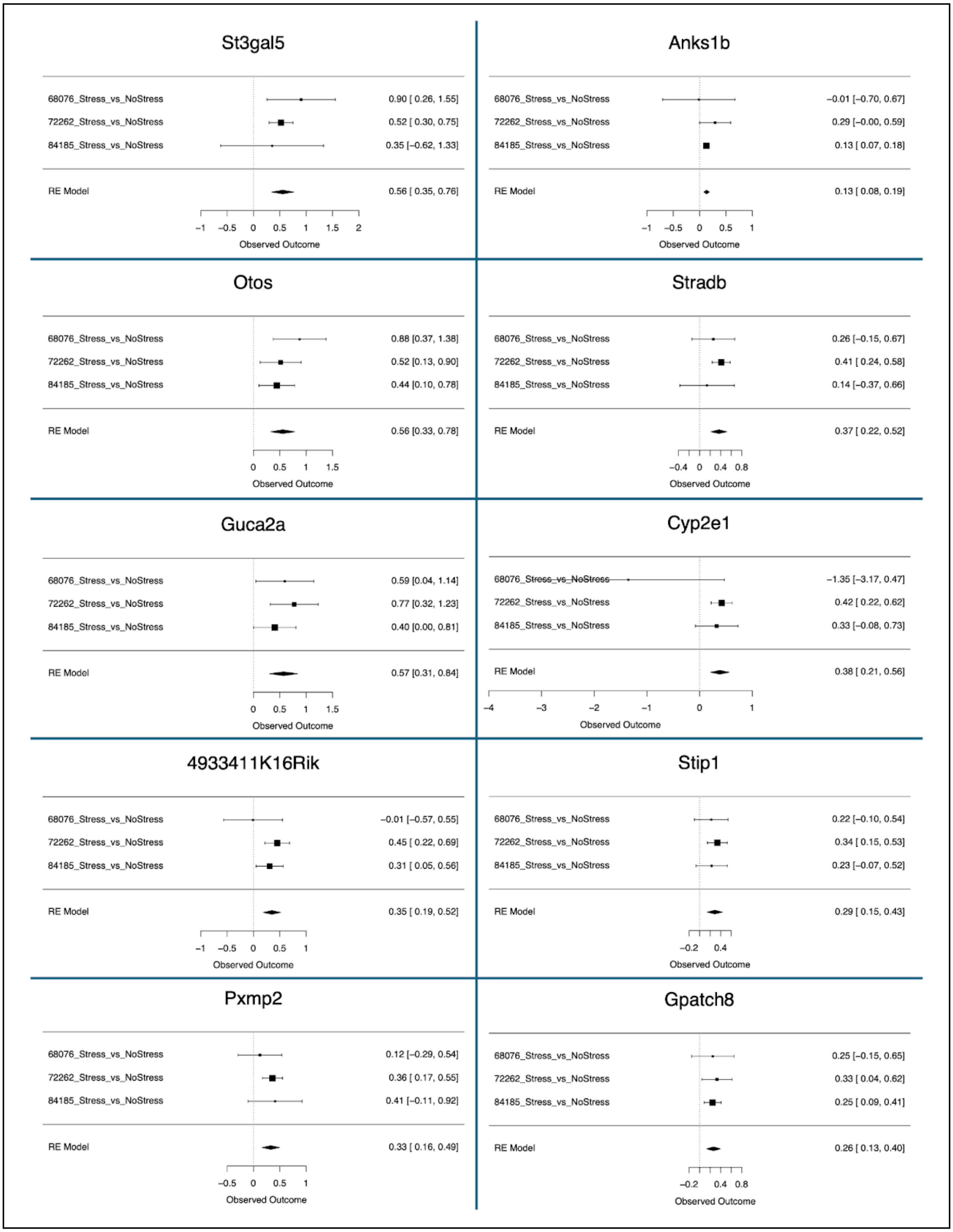

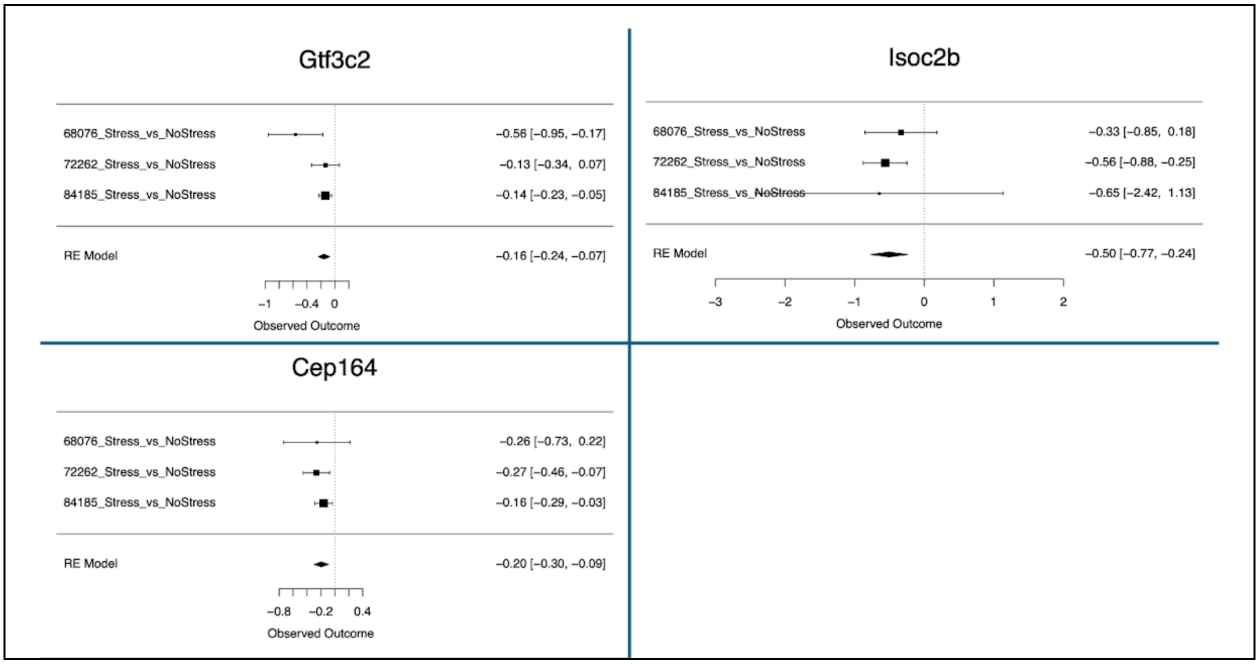


***Figure S1. Forest plots for differentially expressed genes from the meta-analysis (FDR<0.05****)* *Rows illustrate chronic stress Log2FC (squares) with 95% confidence intervals (whiskers) for each of the datasets and the meta-analysis random effects model ("RE Model"). Forest plots allow for visual inspection of the consistency and magnitude of effects across the three studies. Downregulated genes (Figure 2A) have a negative observed outcome for the RE model; Upregulated genes (Figure 2B) have a positive observed outcome for the RE model.*
